# Species-specific phototaxis of coral larvae causes variation in vertical positioning during dispersal

**DOI:** 10.1101/2020.07.31.230235

**Authors:** Aziz J Mulla, Che-Hung Lin, Shunichi Takahashi, Yoko Nozawa

## Abstract

Controlling vertical positioning is a key factor limiting the distance coral larvae can travel, as oceanic currents are faster closer to surface. Currently, the vertical position of coral larvae is assumed to be determined by buoyant, lipid-rich gametes. However, here we show that some, but not all, coral species can control vertical positioning by phototaxis. We first examined the effect of light on the vertical positioning of larvae from five different coral species in the laboratory. We found that larvae from *P. verrucosa*, but not from other coral species, show phototaxis towards light and accumulate near the surface. This behavior was consistent at any age and at any time during the day. In field experiments, using *P. verrucosa* larvae at three different depths (1, 7 and 15 m), the accumulation of larvae in the top half of transparent chambers was observed at all depths. However, such behavior failed to occur in dark chambers. We conclude that larvae from *P. verrucosa*, but not all coral species, accumulate close to the seawater surface as a result of actively swimming towards sunlight. This finding provides a new hypothesis that phototactic behavior is a key factor in regulating vertical positioning for the dispersal of coral larvae.

## Introduction

Reef-building corals are sessile organisms meaning dispersal is limited to the larval stage. Larvae of pioneer coral species can disperse far distances, away from the mother colony to rapidly colonise new areas. Such pioneers usually become the dominant species in areas recently devastated by environmental change i.e. abnormal increases in seawater temperature leading to coral bleaching and mass-mortality. Despite this, reef-building corals remain the cornerstone of coral reef ecosystems, underpinning fundamental components such as structural complexity (Graham and Nash, 2013). Therefore, the ability of larvae to disperse far distances is not only crucial in increasing the distribution of corals worldwide, but also for the establishment of new coral reefs. Free-swimming coral larvae have cilia; however, their dispersal disproportionately relies on oceanic currents. Thus, the ability of coral larvae to influence vertical positioning is a key factor limiting the distance larvae can travel, as oceanic currents are faster closer to the surface.

The regulation of vertical positioning in coral larvae is largely explained by temporal changes in buoyancy, determined by lipid stores (Arai et al., 1993; Rivest et al., 2017). Initial lipid content in coral larvae varies among species; for example, *Acropora palmata* and *Orbicella franksi* have higher lipid contents (~70%) (Wellington and Fitt, 2003) in contrast to *Goniastrea retiformis* and *Stylophora pistillata* (~11–17%) which are significantly lower (Harland et al., 1993). Therefore, some species of coral larvae use lipids to localize close to the seawater surface, taking advantage of the strong currents to disperse, until they lose buoyancy by lipid consumption. Interestingly however, larvae of pioneer coral species *Pocillopora verrucosa*, have noticeably less buoyancy due to extremely low lipid stores (~11%) (Harland et al., 1993). We can therefore consider that buoyancy is not the only factor controlling the vertical position and dispersal of coral larvae.

Kawaguti (1941) observed that four species of coral larvae (*Pocillopora damicornis, Seriatopora hystrix, Galaxea horrescens* and *Euphyllia glabrescens*) showed a positive phototactic response to light, within a certain range of intensity. These observations provided a hypothesis that phototaxis of coral larvae towards sunlight functions to control larval vertical position. Nevertheless, phototaxis of coral larvae has not been reported since. Raimondi and Morse (2000) examined the effect of light, specifically the time of day (midday and dusk), on coral larvae (*Agaricia humilis*), determining that light had little or no effect on swimming behavior. Therefore, the question and significance of phototaxis in coral larvae remains disputed, despite larval behaviour having a potentially critical role in the dispersal process.

In this study we examined whether coral larvae use light to control vertical positioning in five coral species (*P. verrucosa, Pocillopora sp., Dipsastraea speciosa, Favites <u>p</u>entagona*, and *Acropora hyacinthus*). We found that only *P. verrucosa* display a positive phototactic response to light. This response was observed when light was exposed from the top as well as the bottom and side, where larvae accumulated close to the light source. To prove that such light response contributes to the vertical positioning of coral larvae in nature, we also tested this in the field. Our results demonstrate that *P. verrucosa* larvae use light to accumulate near the seawater surface. These results suggest that some, but not all, corals species control vertical position by the ability to sense light signals through the use of phototaxis, providing a new hypothesis that phototaxis of larvae is a factor limiting the ability to localize near the seawater surface and in turn, disperse further.

## Results

In the laboratory, we first examined the effect of light on the vertical positioning of larvae from five different coral species; *P. verrucosa, Pocillopora sp., D. speciosa, F. pentagona* and *A. hyacinthus* (Fig. 1a). Larvae were inserted into black chambers (tubes), placed vertically with or without light directed from the top for 60 minutes. The chambers were then equally partitioned into proximal (P) and distal (D) halves and the distribution index was calculated as [(P-D)/(P+D)]. For *P. verrucosa*, the distribution index was almost 0 after larvae were incubated in darkness, indicating that the vertical position is random. However, under light treatment of 40 μmol m^−2^·s^−1^, the index was close to 0.67, signifying that larvae accumulated at the top of chambers. A similar pattern was observed under a weaker light (0.005 μmol m^−2^·s^−1^) with an index of 0.71. These results show that light influences the vertical position of *P. verrucosa* larvae.

**Fig. 1.**
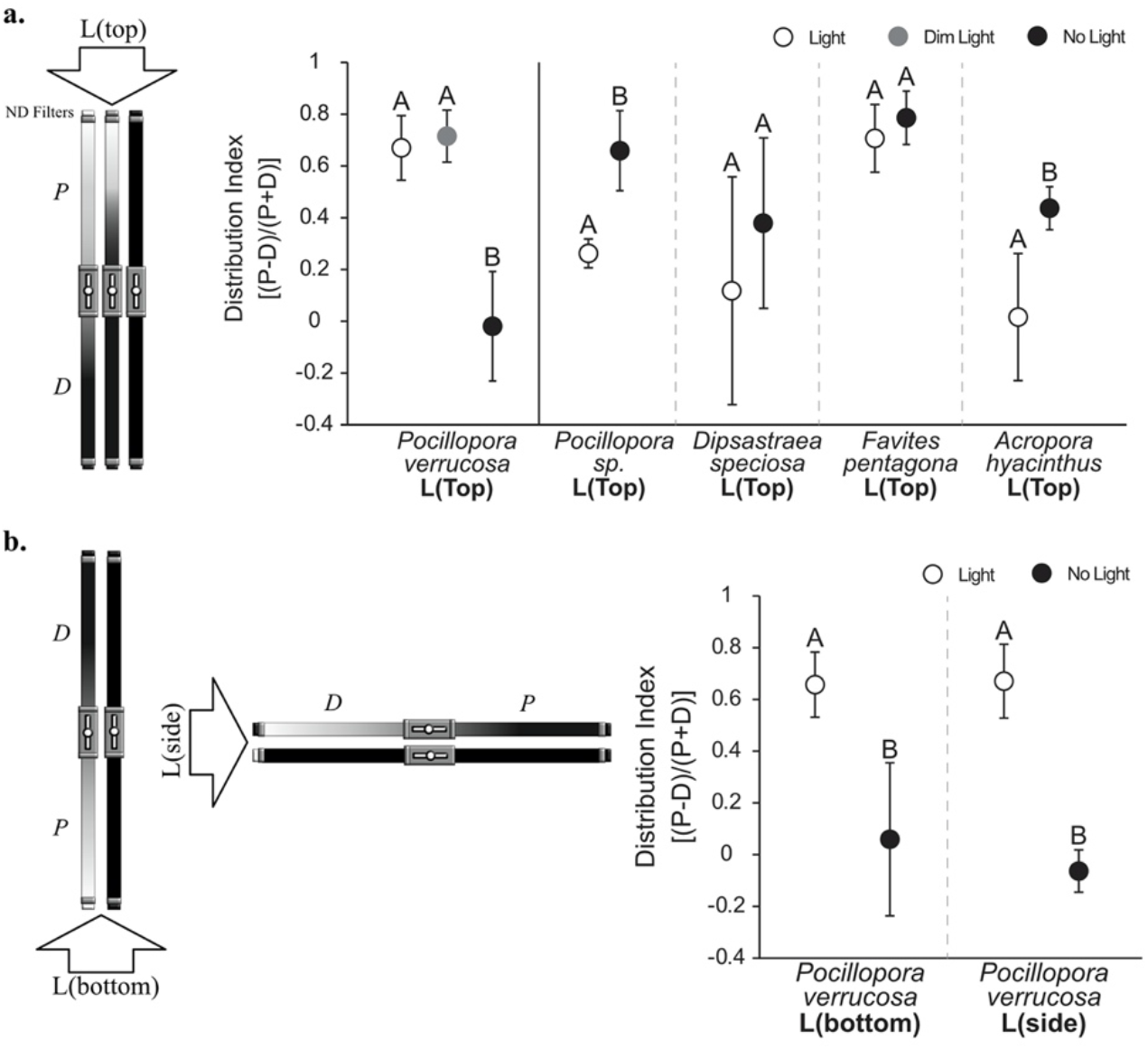
Phototactic response of *Pocillopora verrucosa, Pocillopora sp., Dipsastraea speciosa, Favites pentagona* and *Acropora hyacinthus* in the laboratory. a) Experimental set-up in the laboratory using black chambers (n=15) under a fluorescent lamp using neutral-density filters placed on top of chambers to regulate light intensity. Phototactic response of *P. verrucosa* (n = ~370 per chamber), *Pocillopora sp*. (n = ~40 per chamber), *D. speciosa* (n = ~55 per chamber), *F. pentagona* (n = ~200 per chamber), *A. hyacinthus* (n = ~200 per chamber) to light (white circles; 43.38 μmol m^−2^·s^−1^) and no light (black circles; 0 μmol m^−2^·s^−1^) conditions. All chambers were placed vertically with light positioned from the top. An additional dim light condition (grey circle; 0.0043 μmol m^−2^·s^−1^) was used for *P. verrucosa*. The distribution index indicates the propotion of larvae in the upper and lower sections of chambers, ranging from 1 (all larvae in the upper; positive phototaxsis) to −1 (all larvae in the lower; negative phototaxis), with 0 indicating neutral phototaxis or random swimming (see methods for more detail). Letters *P* and *D* represent proximal and distal respectively. Average ± SD are shown. Letters indicate significant differences (*P* < 0.001) between all active light treatments (light, dim light) and dark treatments (no light) with the exception of *A*. *hyacinthus*, where significant differences were *P* < 0.01. All statistical analyses were conducted within species. b) Phototactic response of *Pocillopora verrucosa* to light from different directions in the laboratory. The light source was provided from the bottom with chambers in a vertical position and then from the side with chambers in a horizontal position. Transparent chambers under bottom and side conditions were split in two (half transparent, half darkened) to observe a clear positive phototactic response. *P*. *verrucosa* larvae (n = ~100 per chamber) were tested under a light treatment similar to that in Fig. 1a (43.38 μmol m^−2^·s^−1^). Average ± SD are shown. Letters indicate significant differences (*P* < 0.001) between light treatment (light) and dark treatment (no light). Statistical analyses were conducted between light treatments.

Of the 4 other coral species, light response varied among larvae, differing from *P. verrucosa* (Fig. 1a). For example, *D. speciosa* and *F. pentagona* larvae did not exhibit any obvious response to light, distributing evenly throughout the chambers with the majority of larvae accumulating in the upper sections, respectively - regardless of light condition. In contrast, the majority of *Pocillopora* sp. and *A. hyacinthus* larvae congregated in the top of chambers in darkness, whereas larvae were observed to distribute more evenly under the light condition (Fig. 1a).

Larval positioning towards the seawater surface is considerable due to a potential phototactic response. Nonetheless, we still cannot exclude that larval movement is regulated by light through photokinesis and motile larvae swim towards the seawater surface via another mechanism i.e. geotaxis. Therefore, we examined whether *P. verrucosa* larvae swim towards light when exposed from the bottom or the side (Fig. 1 b). When light was provided from the bottom of chambers, the distribution index was 0.66, demonstrating that the majority of larvae accumulated near light source (Fig. 1b). This behavior was not seen in darkness where the index was close to 0. Furthermore, a similar result was observed when light was directed from the side (Fig. 1b). These results demonstrate that *P. verrucosa* larvae accumulate near the light source, despite light orientation.

To further understand the effect of light on the vertical positioning of larvae in *P. verrucosa*, larval response to light was measured at different times throughout the day (Fig. 2a) and at different ages (Fig. 2c). Our results demonstrate that larvae have the ability to accumulate close to the surface at any time during the daytime under direct light exposure positioned at the top (Fig. 2a). Under laboratory conditions, a 50% survival rate in *P. verrucosa* larvae was observed at 20 days (Fig. 2b) with a maximum longevity of 24 days. We then compared the effect of light on the vertical position of larvae at 3 and 10 days old. Our results show that there is no difference in larval response to light or the distribution of larvae in chambers as the majority of larvae were in the top under the light condition, but not in darkness (Fig. 2c). These results suggest that light associated accumulation closer to the surface is a persistent response, rather than a temporal one.

**Fig. 2.**
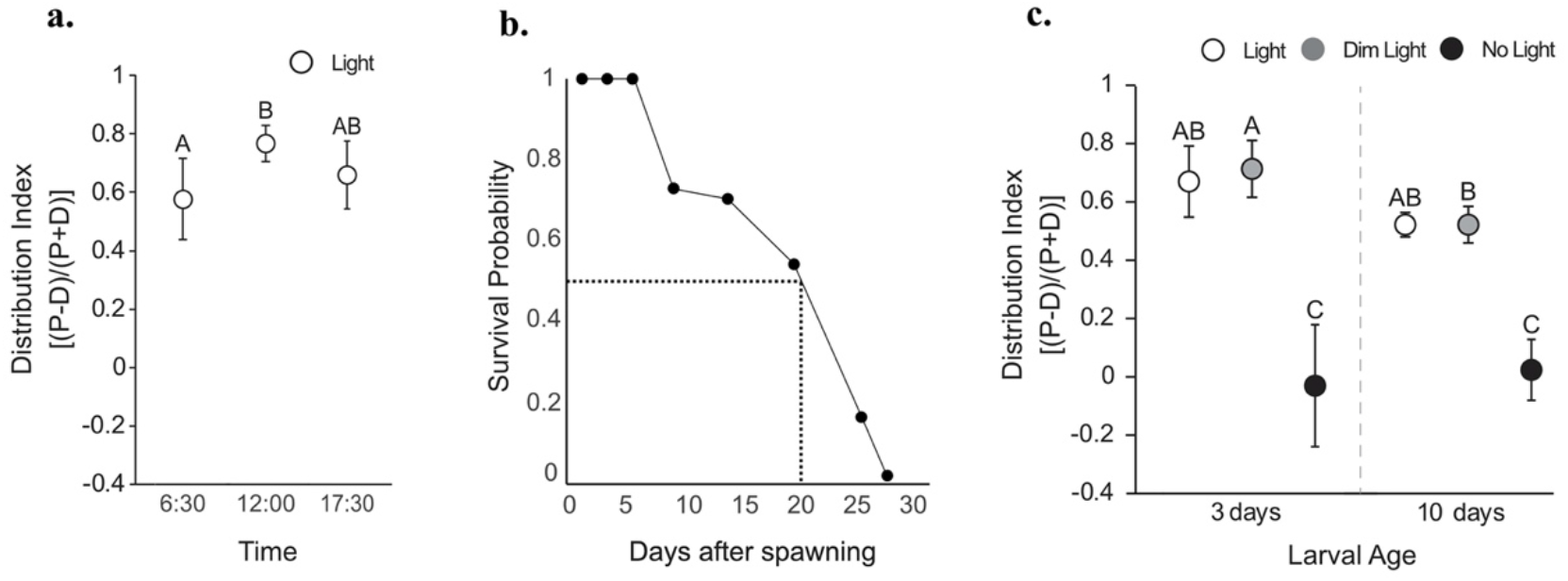
Phototactic behaviour occurs throughout the daytime and when *P. verrucosa* larvae age. a) Phototactic response of *P. verrucosa* larvae tested at various times during sunlight hours. The distribution index is as referred to in Fig. 1. Average ± SD are shown. Letters indicate significant groupings (*P* < 0.05). b) Survival probability of *P. verrucosa* larvae (n=192). c) Phototactic response of *P. verrucosa* larvae at 3 and 10 days old. Larvae at 3 days old (n = ~370 per chamber) and 10 days old (n = ~150 per chamber) were tested under light (43.38 μmol m^−2^·s^−1^), dim light (0.0043 μmol m^−2^·s^−1^) and no light (0 μmol m^−2^·s^−1^) conditions in the laboratory. Average ± SD are shown. Letters indicate significant differences (*P* < 0.001) with the exception of ages under dim light conditions (P < 0.01).

On the basis of our laboratory experiments, we examined larval vertical positioning in response to light in the field in Lyudao (Green Island), Taiwan (Fig. 3). To do so, *P. verrucosa* larvae were inserted into transparent and dark-sealed chambers and placed at three different depths (1 m, 7 m and 15 m) for 60 mins (Fig. 3). The chambers were then equally partitioned into proximal (top) and distal (bottom) halves in the field and the number of larvae in each half was counted to calculate the distribution index. At 1 m depth, the distribution index was 0.56 in transparent chambers and 0.09 in dark-sealed chambers. These results demonstrate that larvae exhibit positive phototaxis towards light in the field. Similar results were observed at 7 and 15 m depths. It is therefore likely that larvae can detect light, both intensity and wavelength, to at least 15 m, resulting in a positive phototactic response.

**Fig. 3.**
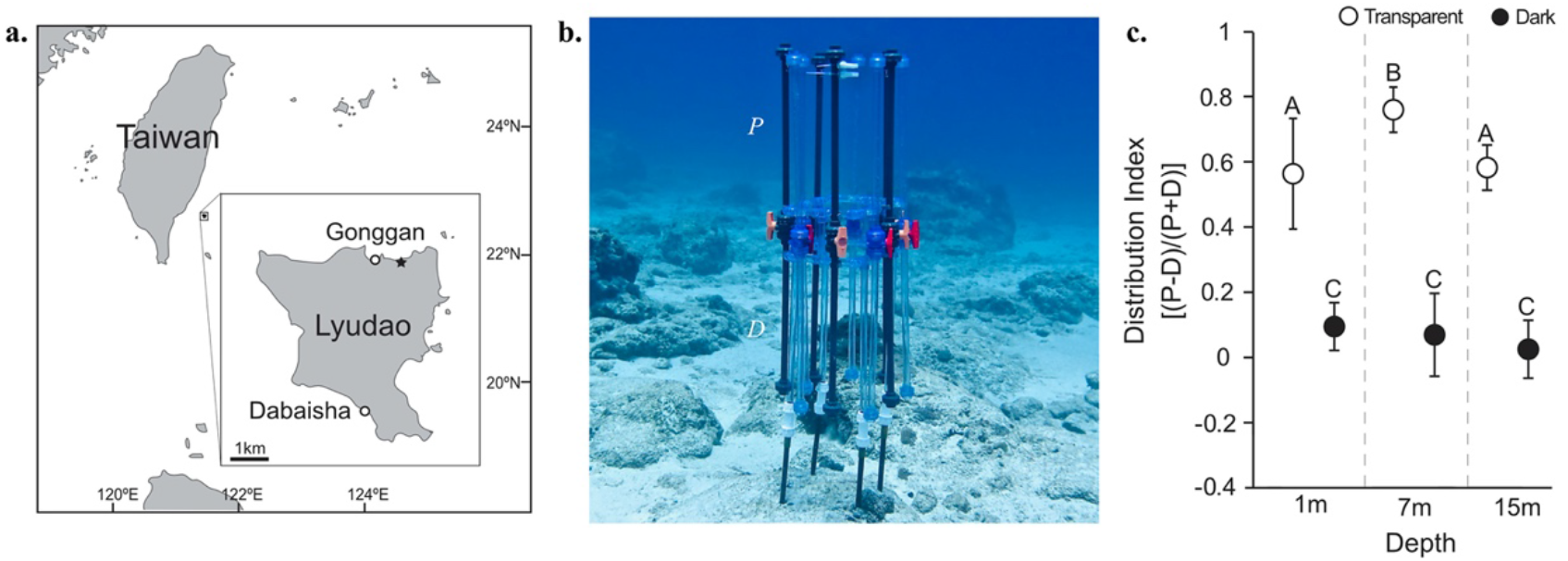
Phototactic response of *Pocillopora verrucosa* larvae in the field. a) Map showing study location within the islands surrounding Taiwan. Daibaisha is where gametes were collected and Gonggan is where experiments were conducted in the field. The star indicates the location of Green Island Marine Station b) Experimental set-up at 15 m showing 5 transparent and 5 black chambers fixed to a frame attached to the reef substrate. c) Phototactic response of *P. verrucosa* larvae (n = ~350 per chamber) *in-situ* at depths of 1 m, 7 m and 15 m in transparent (white circles) and black (dark circles) chambers. The distribution index is as referred to in Fig. 1. Letters *P* and *D* represent proximal and distal respectively. Average ± SD are shown. Letters indicate significant differences between light (transparent) and no light (dark) conditions (A, B vs. C; *P* < 0.001) and within transparent conditions, between depths (A vs. B; *P* < 0.05). Photo credit: A.J. Mulla.

## Discussion

### Larvae of *P. verrucosa*, but not all coral species, accumulate near light stimuli

Kawaguti (1941) reported that larvae of four coral species (*P. damicornis, S. hystrix, G. horrescens* and *E. glabrescens*) swim towards light, indicating that they show signs of positive phototaxis. However, Raimondi and Morse (2000) observed that coral larvae from *A. humilis* larvae showed no significant effect of light on swimming behaviour and positioning. It has therefore remained a controversial issue as to whether or not coral larvae show phototaxis at all. In the present study, we examined the effect of light on the vertical positioning of larvae from five different coral species and found that only larvae from *P. verrucosa* accumulate close to a light source (Fig. 1a). Further, a similar response was seen when *P. verrucosa* larvae were exposed to light from the side or bottom (Fig. 1b). These results suggest that some coral larvae possess phototaxis, but it is not common across all coral species.

Coral larvae swim using cilia that propel, or steer planula in a certain direction (Jékely, 2009). Phototactic steering in larvae can be often observed in marine invertebrates (Angel and Pugh, 2000) and in most cases, steering is regulated by specialized photoreceptors that have photosensory membranes, containing shading pigment granules and cilia that bend upon the direction of light (Leys and Degnan, 2001; Adamska et al., 2007). Animals adjust their entire body orientation in response to a light stimulus, therefore when larvae swim, they move in the direction of the light source. It is uncertain if *P. verrucosa* larvae use this exact mechanism (Harrison and Wallace, 1990), but it has been shown that coral larvae have photoreceptors, such as rhodopsin (Vize, 2009). Furthermore, we are yet to understand why phototaxis is not seen across all coral larvae and what changes occur in order to have such ability. However, as larval motility is common in all coral species used in the study, differences might originate from light reception or signal transduction.

### Light-dependent accumulation of larvae near the seawater surface

The ability to accumulate near the seawater surface is a key factor limiting the distance coral larvae can travel, as oceanic currents are faster closer to the surface. It has previously been shown that larval buoyancy controls vertical positioning (Arai et al., 1993). However, in the present study, we showed, both in the laboratory and field, that larvae from *P. verrucosa* accumulate near the seawater surface in light, but not in darkness. This finding suggests that the vertical positioning of coral larvae can be also controlled by light, through phototaxis (Fig. 4). In general, however, buoyancy of larvae gradually weakens due to lipid loss (Harii et al., 2007). Therefore, the ability to remain close to the surface would be limited in the early larval phase. Conversely, phototaxis of larvae persists throughout the larval stage (Fig. 2). Phototaxis might therefore be a more suitable mechanism to remain at the seawater surface for prolonged durations and as a result, disperse further.

**Fig. 4.**
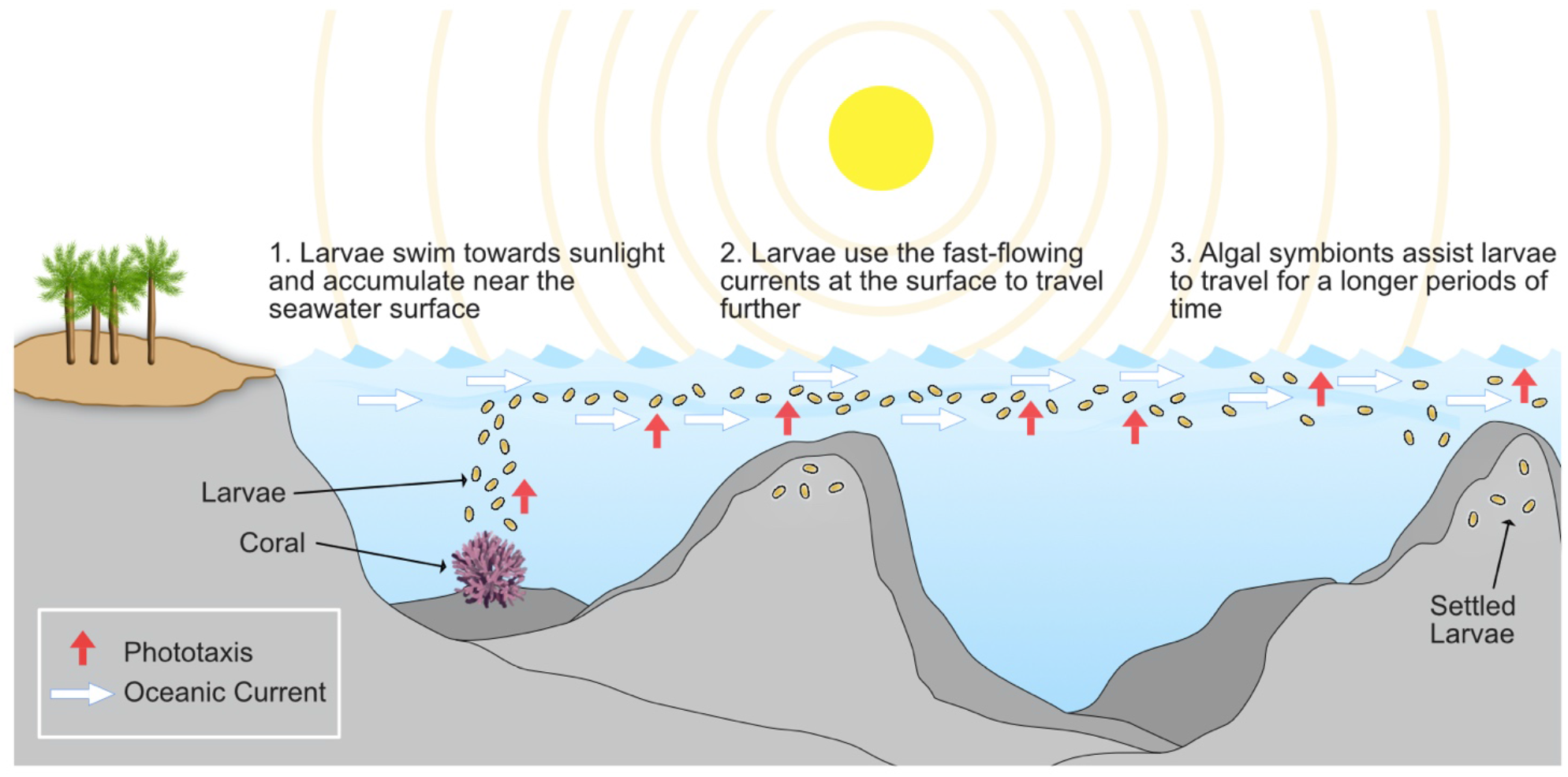
Schematic illustrating the phototactic model proposed for *Pocillopora verrucosa* larvae *in-situ*. Free-swimming larvae fertilized from gametes of *P. verrucosa* swim up towards the seawater surface due to phototaxis (red arrows). Sunlight provides energy for symbionts within the larvae as they dwell at the surface. White arrows represent currents that are faster closer to the surface and larvae use these to disperse farther afield.

Phototaxis in *P. verrucosa* larvae was frequently seen at three different depths in the field, suggesting that this behavior occurs over a wide-range. Furthermore, an accumulation was seen under light at 0.005 μmol m^−2^·s^−1^ in laboratory experiments (Fig. 1a). This result suggests that the light-dependent accumulation of *P. verrucosa* larvae, near the seawater surface, occurs even under the faintest of light, e.g., deep water and cloudy settings. Larvae may also have the ability to respond to moonlight, if they continue to be motile at night. However, further research is required to validate such a hypothesis.

The regulation of vertical positioning by a light signal i.e., phototaxis, has been observed in an array of adult marine organisms such as fish (Wales, 1984), crabs (Shirley and Shirley, 1988), copepods (Kim et al., 2019) and jellyfish (Katsuki and Greenspan, 2013). Similar phenomenon has also been observed in Aiptasia larvae (Foo et al., 2020) and ascidians (McHenry and Strother, 2003). Therefore, control of vertical positioning by phototaxis is not uncommon in marine organisms, as light is a critical environmental factor for life in the ocean. However, our results demonstrate that phototaxis of larvae is species-specific in reef-building corals. Light is therefore not always a factor controlling the vertical positioning of coral larvae, leading to the assumption that coral larvae employ alternative mechanisms to migrate to the seawater surface for dispersal. Since accumulating near the surface is advantageous, in order to travel further via oceanic currents, coral species that are phototactic in the larval phase, might be more suitable for dispersing further (Fig. 4). Consistent with this hypothesis, *P. verrucosa* as a pioneer coral species, has a wide-ranging distribution across the Indo-Pacific, inferring a high dispersal capability.

Despite its obvious dispersal advantage, phototaxis may provide disadvantages for free-swimming larvae, which may explain species-specific responses. As phototaxis causes *P. verrucosa* larvae to actively swim towards light, the continuous consumption of energy could lead to further tradeoffs during the larval phase i.e., longevity. For *P. verrucosa* however, symbionts are transmitted directly from parent to offspring (vertical transmission) (Hartmann et al., 2017). Thus, larvae of *P. verrucosa* are guaranteed to possess symbionts that provide energy for survival (Isomura and Nishihira, 2001). The possession of symbionts might therefore be a key factor for dispersal, especially for those species regulating vertical positioning by phototaxis, as it requires more energy than buoyancy.

Over the past three decades, a myriad of environmental issues have led to mass-mortality events, mainly in the form of coral-bleaching. In the recovery stages of coral reefs, immigration of larvae, via dispersal, is essential to nourish areas devastated by disturbance. In general, some coral species, referred to as pioneers, colonize these areas first. In the present study, pioneer coral species *P. verrucosa* showed that larvae accumulate near the seawater surface via positive phototaxis. This finding suggests that phototaxis might play a key role in the dispersal of *P. verrucosa* as a pioneer species. Accumulation of coral larvae at the surface can be induced by buoyancy, however this is limited to species with high lipid contents. Phototaxis provides an alternative mechanism to reach the surface without the use of buoyancy.

## Materials and Methods

### 1) Study site and species

Our study was conducted during coral spawning season in April and May 2019 (Lin and Nozawa, 2017) in Gonggan, Lyudao (Green Island), Taiwan (22° 40’ 32.61’’ N, 121° 29’ 35.574’’ E). The reef at Gonggan was selected as our study site as it offers a gradual slope to conduct experiments at various depths with ease and is in close proximity to Green Island Marine Station, allowing relatively easy transport of equipment. This area is characterized by a mix of stony corals and rocky reef coral seabed. Both laboratory and field experiments used the larvae of scleractinian coral species *P. verrucosa* (Ellis and Solander, 1786) that inhabits mainly shallow waters, but has been reported at deeper sites in Taiwan (De Palmas et al., 2018; Soto et al., 2018). As a pioneer coral species rapid recovery of *P. verrucosa* has been recorded in Moorea (French Polynesia) over a period of 5 years (Bramanti and Edmunds, 2016). *P. verrucosa* is a hermaphroditic, broadcast spawner that grows rapidly and is abundant across the Indo-Pacific (Brown and Dunne, 1988; Veron, 2000; Ridgeway, Hoegh-Guldberg and Ayre, 2001). Unlike the majority of scleractinian corals, *P*. *verrucosa* spawns during the daytime (Bouwmeester, Berumen and Baird, 2011), 2 to 3 days after the full moon in April and/or May in Taiwan (Lin and Nozawa, 2017). It has been used extensively in a number of physiological studies (Richier et al., 2008; Sawall et al., 2014; Edmunds and Burgess, 2016).

### 2) Gamete collection and larval culturing

*P. verrucosa* spawning occurred between 9:30-10:00am, on the 21^st^ April 2019 at Dabaisha, Lyudao (Green Island), Taiwan. Sperm and eggs were collected by placing plastic bags over branches of 9 spawning colonies allowing gametes to collate inside. Bags were then sealed and transported back to Green Island Marine Station and around 11:15am, were emptied and mixed for cross-fertilization. Upon inspection, eggs were observed to be negatively buoyant. Fertilization occurred within 15 minutes of mixing gametes and cell division was observed. Next, sperm density was reduced by removing the top half of water to prevent bacterial contamination and embryos were transferred into 0.22 μm filtered seawater (FSW). 192 embryos were then removed and placed into individual cells of culture plates with FSW to examine longevity (Fig. 2b). After a period of approximately 10 hours (10:00pm), the first swimming larvae were observed and after 11 hours (11:00pm) the majority of embryos had developed into motile larvae.

### 3) Laboratory Experiments

To test larval reaction to light *ex-situ*, fifteen black chambers were constructed from PVC pipes [top chamber: 47.5 cm in length; bottom chamber: 47.5 cm in length; total length of each chamber: 95 cm (O.D.= 2.2 cm; I.D.=1.7 cm)], separated by a PVC ball valve consisting of two halves; an upper and lower. Five pre-density samples (12 ml) were taken from the larval stock to determine the average number of larvae in 12 ml of seawater (~350 larvae/ 12 ml). Chambers were filled with FSW and larvae were introduced from both ends to allow equal distribution between the top and bottom sections of chambers initially. Chambers were then sealed with either transparent or black caps. The top end of black chambers in the laboratory used transparent caps to allow light inside the chamber.

#### 3.1) Time of Day

We tested *P. verrucosa* larval reaction to light at different times of day using larvae that were 2 days old (6:30, 12:30, 17:30) (Fig 2a). For this, we monitored the reaction to light by placing ~200 larvae in the bottom of chambers (filled with FSW) and after a period of one hour, counted larvae in the top and bottom of chambers.

#### 3.2) Pocillopora verrucosa

*P. verrucosa* larvae used in laboratory experiments were 4 and 5 days old. Once larvae and FSW were inserted into both top and bottom sections of chambers, all fifteen chambers were fixed in place vertically on a frame and placed under a fluorescent lamp (MASTER TL5 HO 54W/865 SLV/40, Philips). In order to regulate light intensity, neutral-density filters were placed on top of chambers. Neutral-density filters reduce the intensity of all wavelengths equally to the desired range (0 μmol m^−2^·s^−1^, 0.0043 μmol m^−2^·s^−1^, 43.38 μmol m^−2^·s^−1^). Light intensities were measured using a light intensity logger (TR-74Ui, T&D, Japan) and then using a spectrometer (LA-105, NK system, Osaka, Japan) to covert lux to μmol m^−2^·s^−1^ through a linear regression model in R (version 3.6.1). As the fluorescent lamp used could alter the surface temperature inside the chambers by creating a gradient, one extra chamber was used to monitor any significant change in temperature of surface water close to the light, recorded at the start and end of the experiment. However, there were no notable changes in temperature over the course of the 1-hour experiment. Valves were opened to allow larvae to swim freely for 1 hour. On collection, valves were sealed then filtered through plankton mesh (100 μm) and larvae from each half of individual chambers was counted under a stereo-microscope.

To assess if larvae moved relative to the direction light (phototaxis) or had increased motility due to changes in light intensity (photokinetic) we set-up two further experiments with light from different directions in the laboratory. In the first experiment, we examined the effect of light from the bottom. We placed 10 chambers in a vertical position, 5 all black chambers and 5 half black and half transparent chambers, with a fluorescent lamp (MASTER TL5 HO 54W/865 SLV/40, Philips) placed at the bottom of chambers. The transparent ends of chambers were facing the light. In the second, we examined the effect of light from the side on horizontal chambers. We placed 10 chambers in a horizontal position, 5 all black chambers and 5 half black and half transparent chambers. A fluorescent lamp (MASTER TL5 HO 54W/865 SLV/40, Philips) was placed at one end of chambers, with the transparent part of chambers facing the light. For both experiments, once an equal number of larvae were inserted into each end of chambers, valves were opened to allow larvae to swim freely for 1 hour. On collection, valves were sealed then filtered through plankton mesh (100 μm) and larvae from each half of individual chambers was counted under a stereo-microscope.

#### 3.3) Other coral species

All experiments for other species (*D. speciosa, Pocillopora* sp., *A. hyacinthus, F. pentagona*) used the same set-up as mentioned in section 3.2. The ages of larvae used for each lab experiments are listed in Table S1.

#### 3.4) Larval survival experiment

To assess the survival curve and approximate longevity of *P. verrucosa* larvae, 192 embryos were removed from the stock and placed into individual cells of culture plates (96-well cell culture plate) with FSW. Larvae were kept in a temperature-regulated growth chamber at 26°C with 12-h exposure to light (06:00–18:00) and dark (18:00–06:00). Every 3 days, FSW was replaced in each cell and mortality was recorded when larvae dissolved.

#### 3.5) Larval age effect experiment

To assess if there was any age-effect on *P. verrucosa* larval reaction to light, we stored the larval stock in a temperature-regulated growth chamber at 26°C with 12-h exposure to light (06:00–18:00) and dark (18:00–06:00). Every 3 days, FSW in the stock was replaced. At 10 days after spawning, we used *P. verrucosa* larvae in the same set-up as described in section 3.2.

### 4) Field Experiments

To test larval reaction to light *in-situ*, thirty chambers of the same design were used (section 3). Fifteen chambers were transparent and fifteen black to block light entirely. All *P. verrucosa* larvae used in field experiments were 5 days old (Table S1). Larvae were collected and prepared as described in sections 2 & 3. Chambers, including larvae, were then transported to Gonggan reef and placed in the appropriate positions on frames located at three depths (1 m, 7 m, 15 m). Data loggers (HOBO Pendant Temperature/Light Data Logger 64K; Onset Computer Corp., Bourne, MA, USA) were placed on both ends of one chamber, at each depth, to monitor temperature and light intensity over time. Valves were opened to allow larvae to swim freely in both sections of the chambers for 1 hour. On collection, valves were sealed, transported back to the marine station where larvae were filtered through plankton mesh (100 μm) and larvae from each half of individual chambers was counted under a stereo-microscope. On the day, the experiment was conducted in the late morning with favourable weather conditions (clear skies, ~30°C). Average sea water temperature and light intensity recorded at each depth is shown in Table S2.

### 5) Distribution Index

The distribution index was determined by using the number of larvae in proximal (P) and distal (D) parts of each chamber, calculated as [(P – D)/(P+D)] (Aihara et al., 2019). Values ranged from −1 to 1 with positive values indicating a positive phototactic response and negative values indicating a negative phototactic response. If the value was 0, neutral phototaxis, or random swimming is implied.

### 6) Statistical Analysis

Due to overdispersion we used beta-binomial generalized linear models to assess any significant differences in the number of larvae in the top of chambers, compared to larvae in the bottom under various conditions (light, dim light, depth, age) in the field and laboratory experiments, as well as to assess any differences between the time of day experiments should take place. We used analysis of deviance (test F) to identify which factors and/or interactions were significant, based on the fitted beta binomial models. Where necessary, we then used Tukey’s post-hoc tests to make pairwise comparisons of values across groups and among interactions. We used statistical software R (version 3.6.1) with package dispmod (version 1.2), package car (version 3.0-5) and package lsmeans (version 2.3) for the analysis.

## Acknowledgments

We thank T-Y Lai, C-L Fong, S. Béniguel, J-H Shiu, M-Y Mok and V. Dang for their assistance in the laboratory. Chiu-Fu Diving Shop and Green Island Marine Station for their field support. This study was funded by an internal research grant of Biodiversity Research Center, Academia Sinica to Y.N. This work was also supported by JSPS KAKENHI Grant Number 20H0330 and 18K19240 to S.T.

## Author Contributions

A.J.M, C.H.L and Y.N conceived the study. A.J.M, C.H.L, S.T, Y.N designed the research. A.J.M, C.H.L and Y.N collected the data. A.J.M analysed the data. All authors wrote the paper.

## Competing Interests

The authors declare no competing interests.

## Notes

### Competing Interest Statement

The authors have declared no competing interest.

