## Supplemental Table 1 and 2 for "Species-specific phototaxis of coral larvae causes variation in vertical positioning during dispersal"

### Tables

**Table S1.** Spawning day and age of larvae used in the experiment.

| Species | Spawning Day | Age of Larvae | | |
|---|---|---|---|---|
| | | *Time of day exp.* | *Lab exp.* | *Field exp.* |
| *Dipsastraea speciosa* | 2019/4/20 | | 4 days old | |
| *Pocillopora verrucosa* | 2019/4/21 | 2 days old | 4 & 5 days old | 5 days old |
| *Pocillopora sp.* | 2019/5/20 | | 4 days old | |
| *Acropora hyacinthus* | 2019/5/23 | | 4 days old | |
| *Favites pentagona* | 2019/5/26 | | 1 day old | |

**Table S2** Average seawater temperature and light intensity at each depth in the field experiment.

| **Depth (m)** | **Temperature (℃)** | **Light Intensity (lux)** |
|---|---|---|
| 1 | 29.2 | 52697.0 |
| 7 | 27.6 | 17378.5 |
| 15 | 27.0 | 506.3 |
